# Embodied decisions as active inference

**DOI:** 10.1101/2024.05.28.596181

**Authors:** Matteo Priorelli, Ivilin Peev Stoianov, Giovanni Pezzulo

## Abstract

Decision-making is often conceptualized as a serial process, during which sensory evidence is accumulated for the choice alternatives until a certain threshold is reached, at which point a decision is made and an action is executed. This *decide-then-act* perspective has successfully explained various facets of perceptual and economic decisions in the laboratory, in which action dynamics are usually irrelevant to the choice. However, living organisms often face another class of decisions – called *embodied decisions* – that require selecting between potential courses of actions to be executed timely in a dynamic environment, e.g., for a lion, deciding which gazelle to chase and how fast to do so. Studies of embodied decisions reveal two aspects of goal-directed behavior in stark contrast to the serial view. First, that decision and action processes can unfold in parallel; second, that action-related components, such as the motor costs associated with selecting a particular choice alternative or required to “change mind” between choice alternatives, exert a feedback effect on the decision taken. Here, we show that these signatures of embodied decisions emerge naturally in active inference – a framework that simultaneously optimizes perception and action, according to the same (free energy minimization) imperative. We show that optimizing embodied choices requires a continuous feedback loop between motor planning (where beliefs about choice alternatives guide action dynamics) and motor inference (where action dynamics finesse beliefs about choice alternatives). Furthermore, our active inference simulations reveal the normative character of embodied decisions in ecological settings – namely, achieving an effective balance between a high accuracy and a low risk of missing valid opportunities.

**Author summary:** In this study, we introduce a novel modeling approach to explore embodied decision-making, where decisions and actions occur simultaneously in dynamic environments. Unlike traditional models that treat decision and action as separate, our framework, based on active inference, reveals that crucial features of embodied decisions – such as feedback loops between decision and action dynamics – emerge naturally. By simulating real-time decision-making tasks, we show how organisms continuously refine their choices by integrating sensory information and motor dynamics. This allows them to strike a balance between decision accuracy and the need for fast, adaptive actions. Our model offers a new perspective on how decisions are influenced by the actions taken, highlighting the importance of considering motor control as an integral part of decision processes. This approach broadens the scope of decision-making research and provides new insights into behavior in ecologically valid, time-sensitive contexts, with potential implications for neuroscience, cognitive science, and fields involving human and animal behavior.

## 1 Introduction

Decision-making is traditionally conceptualized as a serial, *decision-then-act* process, in which sensory evidence is accumulated until a certain threshold, at which point a decision is made and an action is executed. This approach, as formalized by *drift-diffusion* and related models, has been very useful in analyzing behavioral and neural data from laboratory studies of perceptual and economic decisions [1–3]. In these studies, participants select between fixed choice alternatives (usually two) reflecting perceptual judgments (e.g., motion discrimination) or economic offers (e.g., lotteries).

However, animals often face a different class of *embodied decisions*, which imply the choice between courses of actions to be immediately executed in dynamic environments; for example, for a lion, the choice of which gazelle to chase or for a soccer player, the choice of which teammate to which passing the ball [4–10]. To face with the demands of these embodied decisions, animals often need to specify, prepare and sometimes execute actions in parallel to the decision process – as captured by the notion of *affordance competition* [5, 11].

These considerations motivate a series of experiments that use continuous measures of performance during perceptual and economic choices; for example, tracking hand kinematics â using a computer mouse â during the movement from the start position to a response button [12, 13]. Despite their simplicity, these experiments permit analyzing the dynamic processes leading to a decision and the reciprocal influences between the ongoing deliberation and movement (i.e., *decide-while-acting or continuous decisions* [14]). They reveal that participants move and deliberate simultaneously: they generally start moving very early, either toward a specific target or in the middle, if they are more uncertain; they often revisit their decisions in the middle of the trial, as apparent by the curvature of their movements; and they sometimes change mind between the targets, as evident by drastic changes of trajectory [15–18]. These findings eschew serial models and are better explained by parallel [19] or continuous flow models [20] in which unfolding perceptual and decision processes concurrently drive the preparation and possibly the overt execution of one or more responses in parallel – meaning that movements during the task provide a continuous readout of the ongoing deliberation. From a normative perspective, parallel models provide a way to realize decisions faster, which is crucial to survival as it avoids the risk of losing valued opportunity – although sometimes at the cost of reduced accuracy [21].

Crucially, some studies of embodied decisions reveal feedback effects of action dynamics to decision processes that were previously ignored in serial and even parallel decision-making models. For example, a recurring finding is that motor costs associated with different choice alternatives influence perceptual and economic decisions. During ambiguous perceptual decisions [22,23] and value-based decisions [24], participants show a bias to select the response choice associated with the less costly movement. Changes of mind during a perceptual task are less frequent if the costs associated with changing movement direction are greater, such as when response buttons are farther apart [25]. Similarly, during an economic task, changes of mind following a perturbation of the movement trajectory are sensitive to the current state (position and velocity) of the motor system and are less frequent when counteracting the perturbation would be more costly [26].

These and other studies suggest not only that deliberation continues after movement onset (in accordance with parallel models) but also that it is affected by feedback from action dynamics (e.g., motor costs). This motivates a novel class of *embodied decision* models in which action is not the inert outcome of a decision process but influences it, forming a closed loop [21, 27]. These models are motivated by the fact that, from an embodied perspective, the goal of the agent is not just selecting between choice alternatives (as in classical setups) but also simultaneously selecting between potential courses of action to reach the targets (often within a deadline) and tracking the action itself – which means that both decision and action processes need to be jointly and continuously optimized. In turn, at the neural level, embodied decisions might require a *distributed consensus* across various brain networks that process outcome values and motor plans, rather than a centralized process as traditionally assumed [28].

Here, we show that the key signatures of embodied decisions emerge naturally in *active inference*, a framework that models perception and action selection as two aspects of the same objective of free energy minimization [29–33]. By simulating embodied decisions as an active inference process, we are able to reproduce various empirical findings about the parallel unfolding of actions and decisions in time, as well as feedback effects of movement dynamics in perception. Furthermore, we illustrate the normative advantages of embodied choices over serial choices under time pressure.

## 2 Results

We illustrate the functioning of an active inference model of embodied decisions by simulating a two-alternative forced choice (2AFC) decision task with time-varying information, i.e., in which evidence for one choice or the other, expressed in terms of sequentially provided cues, changes throughout each trial, as in [34, 35] (Fig 1a). The agent has to move the 3-DoF arm from a start position (small blue dot at the center) to reach a left (red circle) or right (green circle) target button, to report which target has or will have more cues in it. During the task, 15 cues appear, one after the other, either in the left or the right circle, and then disappear leaving only the last one visible. The agent can start moving at any moment and the cues continue to appear normally during movement. The trial ends when the agent reaches one of the two buttons (or within a deadline).

**Fig 1.**
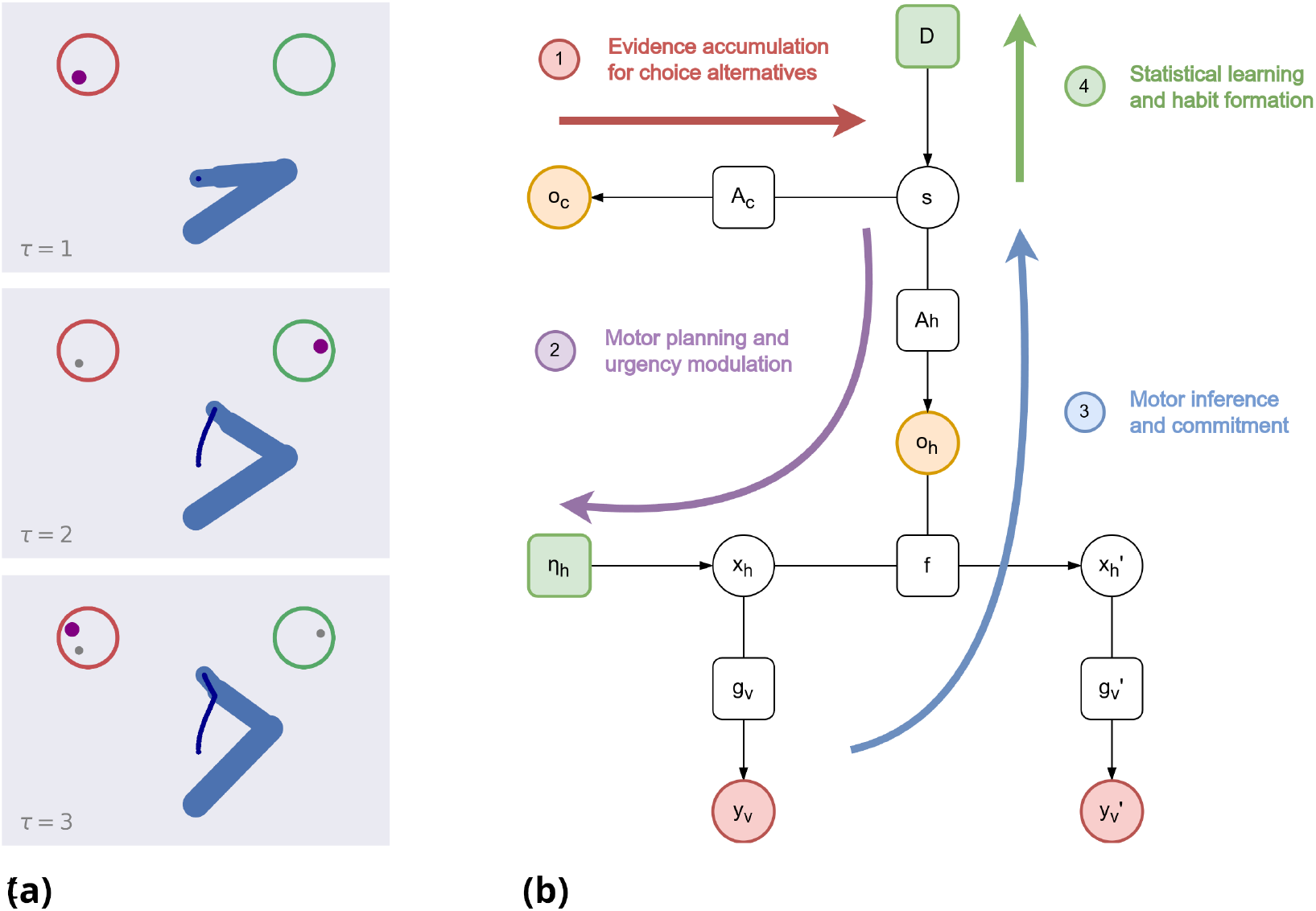
Embodied decision setup and active inference model. (a) Experimental setup, during three consecutive discrete time steps *τ*. The agent controls a 3-DoF arm (the three segments in blue), which starts at a home position (blue dot) at an equal distance from the two targets (red and green circles). The current cue is represented with a big purple dot, while the old cues are represented with smaller grey dots. For each trial, the agent has to reach with the hand the target it believes will contain more cues. The hand trajectory is represented with a thinner blue line. (b) Hybrid active inference model for embodied decisions. The model comprises four processes, numbered from 1 to 4. In the first process, discrete hidden states ***s***, encoding the probability that each target is the correct choice for the current trial, are iteratively inferred by discrete cues ***o***_*c*_ by inverting the cue likelihood matrix ***A***_*c*_. In the second process, the hidden states ***s*** generate a particular combination of discrete hand dynamics ***o***_*h*_ through the extrinsic likelihood matrix ***A***_*h*_. Each hand dynamics *o*_*h,m*_ is related to a continuous dynamics function ***f***_*m*_, where the target positions are defined (see [36, 37] for more details). A forward message imposes a prior over the hand velocity (in Cartesian coordinates) 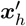, while a backward message infers the related Cartesian position of the hand ***x***_*h*_, ready for kinematic and dynamic inversions. In the third process, for each continuous time step *t*, the current position and velocity of the hand (in a visual domain) are inferred by continuous observations ***y***_*v*_ and 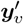, via the corresponding likelihood functions (***g***_*v*_ and 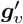). The inferred hand trajectory flows back toward the prior ***η***_*h*_ for action execution (see Section 4.2). For each discrete time step *τ*, the target probabilities ***s*** are also inferred by the current motor trajectory. Finally, in the fourth process, the prior ***D*** over the correct target is updated across trials, implementing habitual learning. Of note, some readers may find the edge from ***s*** (the agent’s belief about the latent state) to ***o***_*c*_ (an observation) somewhat unconventional. This is because, in many contexts, observations are typically assumed to be generated from the true latent states rather than the agent’s beliefs about them. However, this formulation is standard in active inference studies, even when representing the agent’s generative model as in this figure.

Crucially, by manipulating the sequence of cues, we compare the agent’s decision dynamics in three conditions (or trial types): *congruent*, in which a greater proportion of cues initially appears in the correct target; *incongruent*, in which a greater proportion of cues initially appears in the incorrect target; and *neutral*, in which the proportion of cues initially appearing in both targets is balanced.

Empirical evidence indicates that these trial types significantly influence choice and movement dynamics. In the Tokens task [34], which is conceptually similar to this task (except that the old cues disappear), trial incongruence determines a greater number of errors and longer response times [34] and impacts the lateral hand position prior to a decision, when movement kinematics were recorded [38]. Similar results are observed in the Eriksen flanker task [35], which shares conceptual similarities with our task, because a correct visual target is presented along with congruent or incongruent cues. The flanker task can be modeled as a progressive shift of attention toward the correct target over time, equivalent to sampling it from increasingly precise distributions [16]. This process explains the significant decrease in performance during incongruent conditions (*flanker effect*), because the initial sampling process is biased toward the incorrect response. The decrease in performance manifests itself as increased number of errors, reaction time, and curvature of the movement trajectory in tasks that tracked movement kinematics [16, 39]. Furthermore, measures of motor potentials (ERPs: event-related brain potentials and EMG: electromyographic activity) after target presentation suggested that competing responses were activated simultaneously [40], in keeping with analogous findings through single-cell recordings of the monkey dorsal premotor cortex [41].

Below, we show that a hybrid active inference model (i.e. a model composed of both discrete and continuous variables) that jointly optimizes decisions and actions reproduces these signatures of embodied choices. The model can be decomposed into four interacting processes – evidence accumulation, motor planning, motor inference, and statistical learning across trials (along with habit formation) – see Fig 1b for a schematic illustration and Section 4.1 for technical details. Below, we discuss these processes and present simulations about how they affect the agent’s decision and action processes.

### 2.1 The first process: evidence accumulation for the choice alternatives

The first process is responsible for the accumulation of sequential evidence for the choice alternatives. It includes discrete hidden states ***s***, which encode the probability that each target is the correct choice for the current trial (i.e., the one that will contain the most cues). They are sampled from a categorical (here, binomial) distribution, i.e., ***s*** = *Cat*(***D***) = [*s*_*t*1_ *s*_*t*2]_, where ***D*** are the parameters of a Dirichlet distribution and define the agent’s prior beliefs. In the following simulations, we initialize them with a uniform distribution for each trial. The discrete hidden states ***s*** generate two discrete predictions in parallel. The former – computed through the likelihood matrix ***A***_*c*_ – is a prediction of the cue that the agent will observe next, with *α*_*c*_ playing the role of a precision (i.e., a confidence or certainty in the likelihood mapping between cause and consequence), similar to the drift rate in drift diffusion models. The latter – computed through the likelihood matrix ***A***_*h*_ – corresponds to the hand dynamics: it predicts whether the hand will move toward the left target, toward the right target, or not move at all (with probability 1 − *α*_*h*_). In short, precision or inverse uncertainty (i.e., ambiguity) of the two mappings affects how fast evidence accumulation unfolds, and which movement strategy the agent adopts:

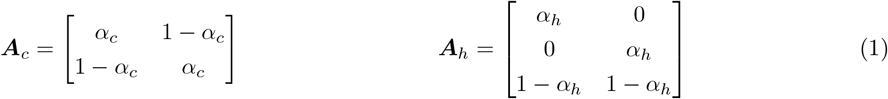

Since the agent has to update its (Bayesian) beliefs based upon continuous observations, the requisite probability transition matrices ***B*** (as defined in Section 4.1) reduce to the identity matrix; such that in the absence of any observations, posterior estimates of latent or hidden states do not change. At each discrete step *τ*, a particular cue ***o***_*c*_ and a particular hand dynamics ***o***_*h*_ are observed and compared with the corresponding predictions. Hence, the inference of the discrete hidden states ***s*** follows the equation:

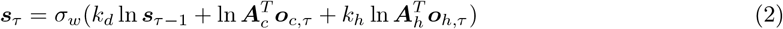

where *σ*_*w*_ is a weighted softmax function:

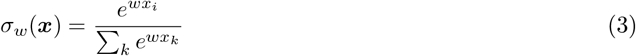

with slope (or precision) *w*. A high value of *w* ensures fast transitions between discrete states, thus avoiding positions within the two targets. In short, the discrete update is a combination of a prior from the previous step (which is equal to ***D*** at the beginning of the trial) and two likelihoods. The first contributes to the accumulation of sensory evidence and iteratively refines the choice based on the sensory cues. The second likelihood links target estimation and hand dynamics; in this way, the variable ***o***_*h*_ behaves as a sensory signal for the discrete model (similar to ***o***_*c*_) and permits accumulating evidence from the agent’s movements. We will unpack the role of the second likelihood in the next sections; here, to simulate the standard evidence accumulation, we let the third term depend on a parameter *k*_*h*_, which we set here to 0 so that the inference of the correct choice only relies on sensory cues ***o***_*c*_. Finally, we include a parameter *k*_*d*_ acting as a forgetting factor (which might be useful to deal with non-stationary tasks), which we keep fixed at 1 in our simulations. Also, we set *α*_*h*_ = 1.0 (so that the agent starts moving at the beginning of the trial) and *α*_*c*_ = 0.6.

We test the active inference model with the three conditions explained above. In *congruent* trials, cues move toward the correct target with an initial probability of 80%, which then gradually increases and reaches 100% after 8 cues. In *neutral* and *incongruent* trials, the probabilities of the correct target are respectively initialized to 50% and 20% and then increase as in congruent trials. Each trial comprises 21 discrete time steps *τ*, each in turn comprising 30 continuous steps *t*. At *τ* = 0, no cue is presented, but the agent can move. For the next 15 time steps, a cue per time step is presented. Finally, in the last 4 time steps, no cue is presented, but the agent can still move and reach the target.

Fig 2 shows that, under all conditions, when movement onset is immediate the agent moves in between the two targets. In the *incongruent* condition, the agent first moves toward the wrong target and eventually changes mind. These results match qualitatively key empirical findings discussed in the Introduction [15–18]. To assess whether the model generates statistically different trajectories under the three conditions, we simulated 100 trials per condition and considered a widely used index of choice uncertainty: the *maximum deviation* of the trajectories from an ideal, straight line between the start and the correct target [10, 12]. We found significantly larger maximum deviation in *incongruent* (M = 125.97, SD = 25.0) compared to *neutral* (M = 88.19, SD = 30.73) trials and *neutral* compared to *congruent* (M = 56.64, SD = 18.56) trials (for all tests, *p* < .001).

**Fig 2.**
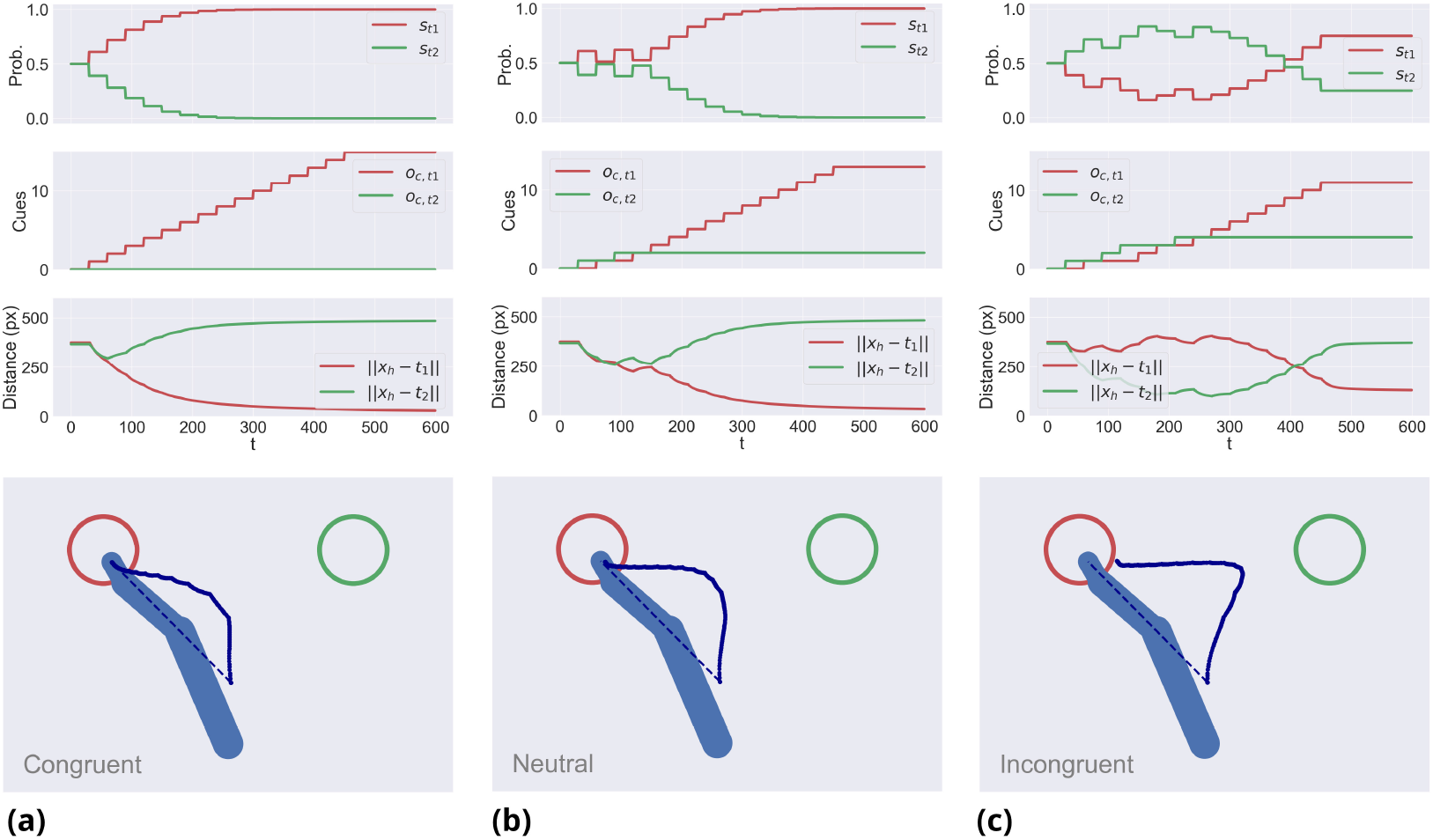
Evidence accumulation during (a) congruent trials, (b) neutral trials, and (c) incongruent trials. The first row shows the dynamics of a sample trial for each condition: specifically, the first plot shows the discrete hidden states ***s*** encoding the two target probabilities over continuous time; the second plot shows the cumulative sum of the cue observations ***o***_*c*_; the third plot shows the distances between the hand and the two targets. Note that the discrete signals are maintained for a whole discrete period *τ*, generating a stepped behavior. The second row shows the agent’s average trajectory (in dark blue) across 100 trials for each condition. A dotted line of minimum distance between the initial hand position and the left target is also displayed.

### 2.2 The second process: motor planning and urgency

When facing the same task, different groups of participants might show different strategies; for example, a conservative strategy to postpone movement until they feel sufficiently confident, or a risky strategy to guess the correct choice and start moving immediately [42].

In our model, the selection of a conservative versus risky strategy depends on two things. First, the rate of evidence accumulation depends upon the precision parameter *α*_*c*_, such that a greater precision (i.e., less ambiguity) enables the agent to form more precise posterior beliefs about the correct target. Second, the movement urgency rests upon the precision parameter *α*_*h*_ (of the likelihood mapping to kinematics), which determines the confidence – the agent has about the correct target – required to initiate movement. Recall that ***o***_*h*_ (the *discrete set of hand dynamics*) encodes the probability that the two targets generate a movement toward the left, toward the right, or no movement (i.e., ***o***_*h*_ = [*o*_*h,t*1_, *o*_*h,t*2_, *o*_*h,s*_]).

Importantly, ***o***_*h*_ specifies potential action plans, as it communicates with continuous hidden states specifying the instantaneous trajectory (position ***x***_*h*_ and velocity 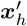) of the agent’s hand in extrinsic (e.g., Cartesian) coordinates. Specifically, each element of ***o***_*h*_ is linked to a dynamics function:

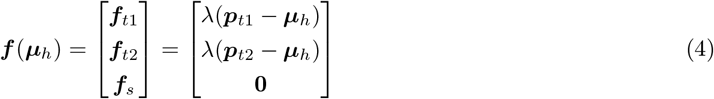

where ***µ***_*h*_ is the belief over the hand position ***x***_*h*_, *λ* is an attractor gain, while ***p***_*t*1_ and ***p***_*t*2_ are the positions of the two targets – assumed to be known and fixed. This mechanism implements the simultaneous preparation of competing motor plans, as also reported in monkey premotor cortex [41].

The mapping between discrete and continuous signals in these hybrid models — with mixed continuous and discrete states — rests on a form of Bayesian model averaging; namely, estimating the movement trajectory by averaging over some discrete models (i.e., hidden states) generating possible trajectories – as explained in Section 4.1. In particular, a prior 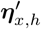 over the hand velocity 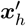 is computed by weighting the three potential trajectories (to reach the two targets and to stay) with their respective probabilities ***o***_*h*_ that the agent plans for a given discrete goal:

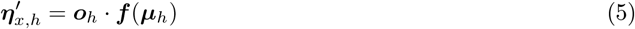

This desired velocity enters the update of the continuous hidden states as a dynamics prediction error 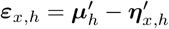 cexpressing a composite motion that the continuous model will realize. The inverse process, i.e., the inference of the current trajectory, is explained in the following section, and more details are found in Section 4.1.

Fig 3 illustrates the effect on movement onset and velocity with three levels of urgency, during an incongruent trial. Since *k*_*h*_ = 0, the evidence accumulation is the same for the three cases, but the trajectories change depending on the agent’s urgency to move, giving rise to risky, medium and conservative strategies. High and intermediate levels of urgency produce riskier strategies that initially move toward the wrong target and then manifest changes of mind. Low urgency produces a conservative strategy that moves directly toward the correct target, but has higher reaction times when the two target probabilities are too close, failing to complete the trial within the deadline. This is because the trajectories generated by the discrete model are constantly weighted by the *stay* dynamics, hence low precision *α*_*h*_ (which means low urgency) results not just in late movement onset but also in slower motion. This simulation illustrates that manipulating urgency provides flexibility in the link between evidence accumulation and movement dynamics. With high urgency, the agent moves earlier and takes the risk of failing, whereas with low urgency, the agent may wait until it accumulates sufficient evidence to reach very high confidence about the correct target. Interestingly, urgency and speed of evidence accumulation can interact, as shown in Fig 4. Finally, this example illustrates that when urgency is set to a very low level, the embodied model approximates – or even transforms into – a serial *decide-then-act* model, initiating movement only after complete evidence accumulation. This result highlights the critical role of urgency in shaping the interaction between decision-making and movement.

**Fig 3.**
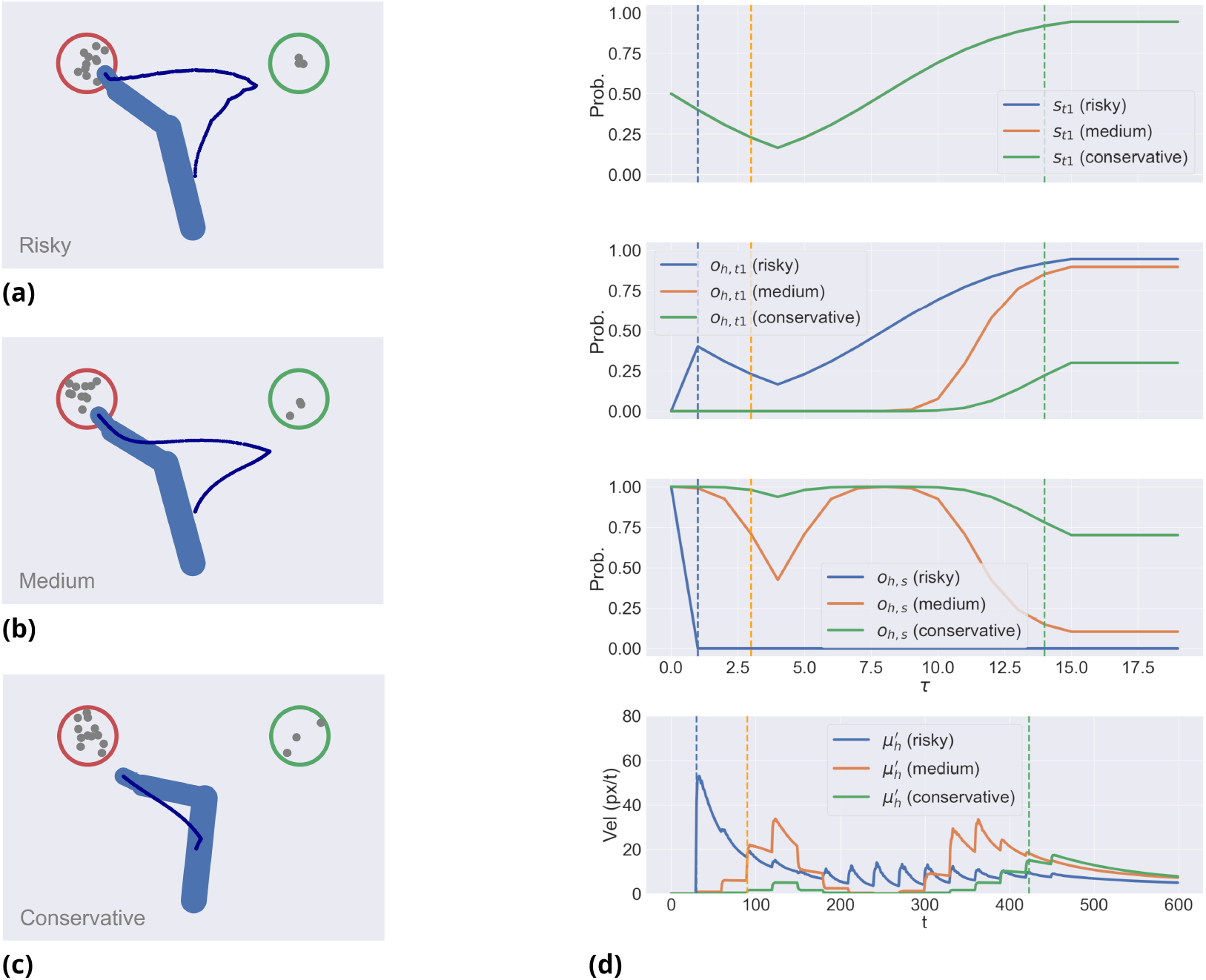
Motor planning with (a) a risky strategy (high urgency, or *α*_*h*_ = 1.0), (b) a medium strategy (medium urgency, or *α*_*h*_ = 0.55), and (c) a conservative strategy (low urgency, or *α*_*h*_ = 0.5). (d) First panel: dynamics of the discrete hidden state of the first target over discrete time *τ*, which in this case are the same for every strategy. Second panel: dynamics of the discrete variable *o*_*t*1_ of the first target. Third panel: dynamics of the discrete variable *o*_*s*_ of staying in position. The fourth panel shows the L2-norm of the belief 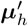 over the hand velocity in continuous time *t*. The vertical dashed lines represent the movement onset for each strategy (using 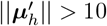 as threshold).

**Fig 4.**
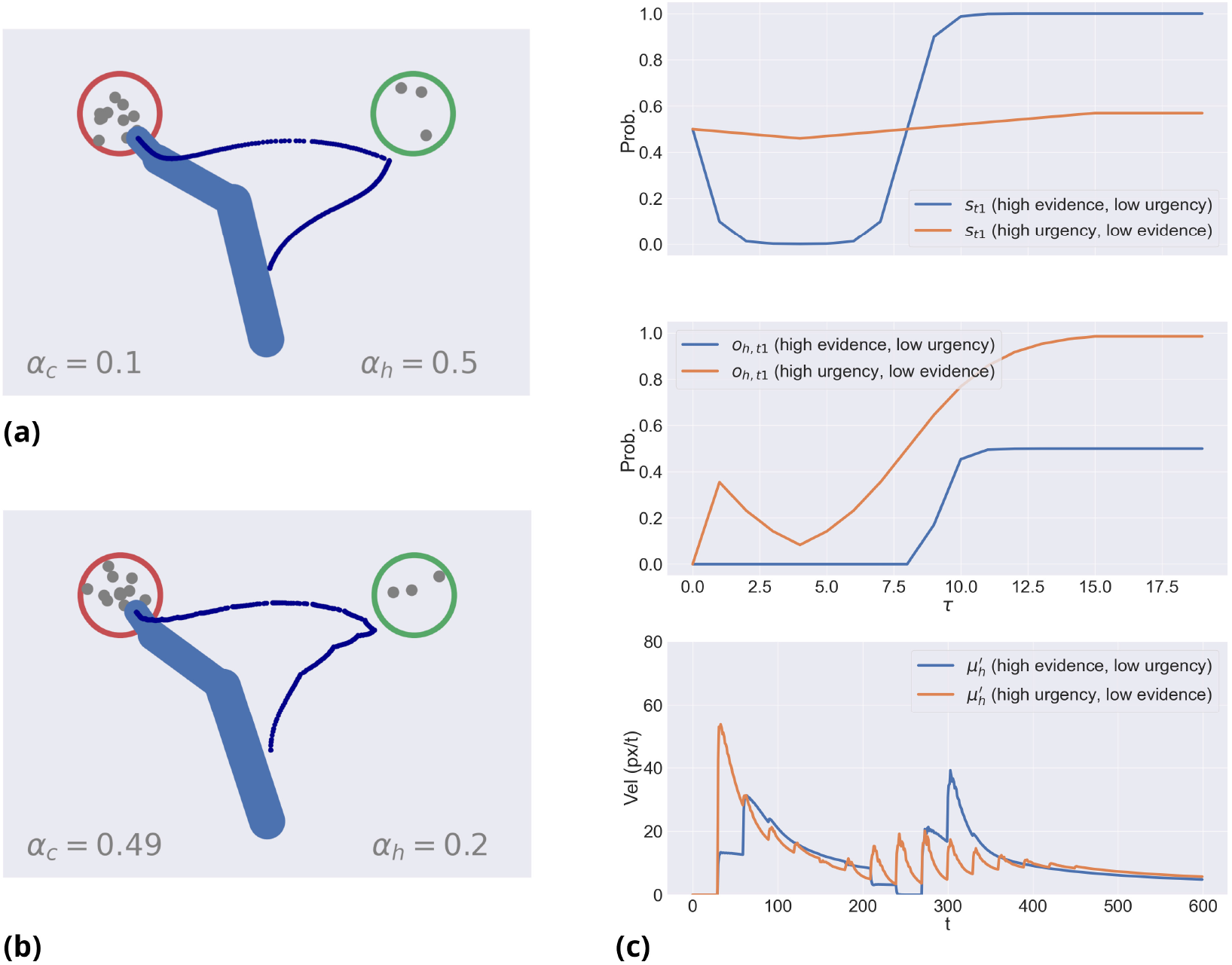
Interaction between urgency and speed of evidence accumulation. (a) Fast evidence accumulation (*α*_*c*_ = 0.9) and low urgency to move (*α*_*h*_ = 0.5). (b) Slow evidence accumulation (*α*_*c*_ = 0.51) and high urgency to move (*α*_*h*_ = 0.8). (c) While movement dynamics look similar, the evolution of the hidden states and hand dynamics are different.

### 2.3 The third process: motor inference and commitment

Various studies found that participants who initiate a movement show high commitment to the initially selected target – even in the face of contrasting evidence – when the costs required to reach the alternative target increase [25, 26]. Interestingly, in our model, this commitment emerges automatically during model inversion. As the agent jointly infers the correct target and the optimal discrete hand dynamics to reach it, there is a reciprocal interaction between the (top-down) process of *motor planning* and its dual (bottom-up) process of *motor inference* (Fig 1b). This is because in our model, not only the agent makes predictions over the cue that will be observed next (i.e., ***A***_*c*_***s***), but also over the hand trajectory (i.e., ***A***_*h*_***s***). The latter prediction entails a causal relation between discrete goals (in this case, the probability of the two targets) and the agent’s movements (here, the dynamics needed to reach the targets) – as explained in Section 4.1. The key implication of this perspective is that the probability of the two targets can be estimated from the hand trajectory itself (***o***_*h*_). As a consequence, a *self-evidencing* mechanism takes place, during which a target inferred by some cues produces a plan to reach it, which in turn confirms the initial agent’s estimate. In other words, via motor inference, movement *stabilizes* (i.e., reduces the uncertainty about) the decision and creates commitment to the initially selected target.

The rich interplay between evidence accumulation, motor planning and motor inference gives rise to three competitive processes. The first competition occurs when estimating the continuous hidden states of the hand trajectory, defined by the generalized beliefs 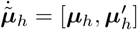:

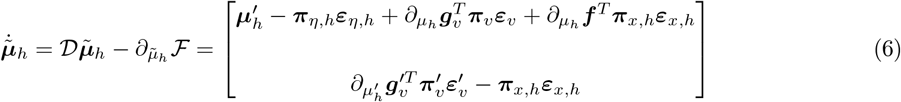

This update rule comprises three main components. First, a prior prediction error ***ε***_*η,h*_ (along with its precision ***π***_*η,h*_) that biases the belief over the hand position. Although not shown in the model, the latter is linked, through forward kinematics, to an intrinsic continuous model encoding proprioceptive trajectories (e.g., expressed in joint angles). As a consequence, the backward message sent from the belief over the hand position to the intrinsic model performs inverse kinematics, eventually driving action. See Section 4.2 and [43, 44] for more details about kinematic inference. Second, visual prediction errors ***ε***_*v*_ = ***g***_*v*_(***µ***_*h*_) − ***y***_*v*_ and 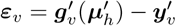 computed through likelihood functions ***g***_*v*_ and 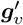, and expressing the difference between predicted and observed hand position/velocity. These terms are backpropagated for both orders, to keep the belief close to the actual hand trajectory. Third, the dynamics prediction error ***ε***_*x,h*_ defined in the previous section, affecting both orders as a forward or backward message.

The second (and most interesting) competition happens when this continuous belief clashes with the desired hand dynamics needed to reach the selected target. While from a top-down perspective the dynamics defined in Eq 4 act as potential trajectories averaged by the probabilities ***o***_*h*_, from a bottom-up perspective they are used to infer the most likely explanation of the real trajectory. More formally, the discrete hand dynamics ***o***_*h*_ at time *τ* is found via Bayesian model comparison, i.e., by comparing the discrete prediction ***A***_*h*_***s***_*τ*_ with the continuous evidence ***ℒ***_*h*_ of the hand trajectory, accumulated over a time window *T* :

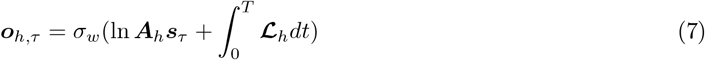

As before, *σ*_*w*_ is a weighted softmax whose slope *w* here controls how fast the transition between different dynamics occurs. High and low values of *w* correspond to abrupt and gradual movement onsets, respectively. In practice, evidence accumulation reduces to a sum over the continuous steps that comprise a single discrete step. For each *m*th dynamics of Eq 4, the log evidence ℒ_*h,m*_ is computed by:

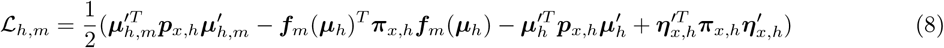

More details about the reduced posteriors 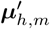 and prior precisions ***p***_*x,h*_ can be found in Section 4.1, and in [36, 37, 45]. Here, we note that each potential dynamics function ***f***_*m*_ is compared to the current dynamics ***µ***_*h*_ inferred via sensory observations. As a result, the log evidence ***ℒ***_*h*_ assigns higher values to the potential trajectories that better match the real one. For instance, if the arm moves toward the red target, the difference in magnitude between the real trajectory and the potential trajectory associated with reaching the red target will be lower, while the one associated with the opposite trajectory will be higher. Hence, the agent can understand which of the two targets is reaching. This evidence is further compared with the discrete prediction ***A***_*h*_***s***_*τ*_ encoding the agent’s desires (reaching the red target, reaching the green target, or staying), so the final hand dynamics will be a combination of the two contributions.

The third and final competition emerges at the intersection between motor inference and evidence accumulation, as highlighted in Eq 2. Since the discrete hand dynamics are constantly steered toward the current hand trajectory, the latter becomes a predictor for the correct target. This ultimately produces a commitment toward the chosen dynamics – expressed in the rightmost term of Eq 2.

One way to appreciate commitment is to consider that, as the distance between the two targets increases, human participants make fewer changes of mind [25]. In line with this, when simulating 100 *neutral* trials with targets at different distances, we found that changes of mind were more frequent with targets at low distance (*n* = 29) than at medium (*n* = 20) and high distance (*n* = 8) – see Fig 5 for some sample trials. The reason for this result lies in Eq 7. Since the discrete hand dynamics ***o***_*h*_ are used to infer the correct choice alternative, closer targets are scored with greater probabilities than farther targets. Furthermore, since the softmax function amplifies the differences between current and potential trajectories, a target is assigned an increasingly lower probability, the farther it is from the other one. This is evident in Fig 6, which shows the potential trajectories to reach both targets for the entire trial, the actual trajectory, and the log evidence accumulated over time. Another way to appreciate commitment is by comparing the agent’s behavior during *congruent* and *incongruent* conditions, with motor inference (*k*_*h*_ *>* 0) and without it (*k*_*h*_ = 0) – see Fig 7. When motor inference is active in the *congruent* condition (Fig 7b), the hand dynamics stabilize the decision: the target is inferred faster than without motor inference (Fig 7a) and the second wrong cue (observed at *τ* = 9) is ignored. When motor inference is active under the *incongruent* condition (Fig 7d), the agent could commit to a wrong decision ignoring contrasting evidence – a behavior that is not observed without motor inference (Fig 7c).

**Fig 5.**
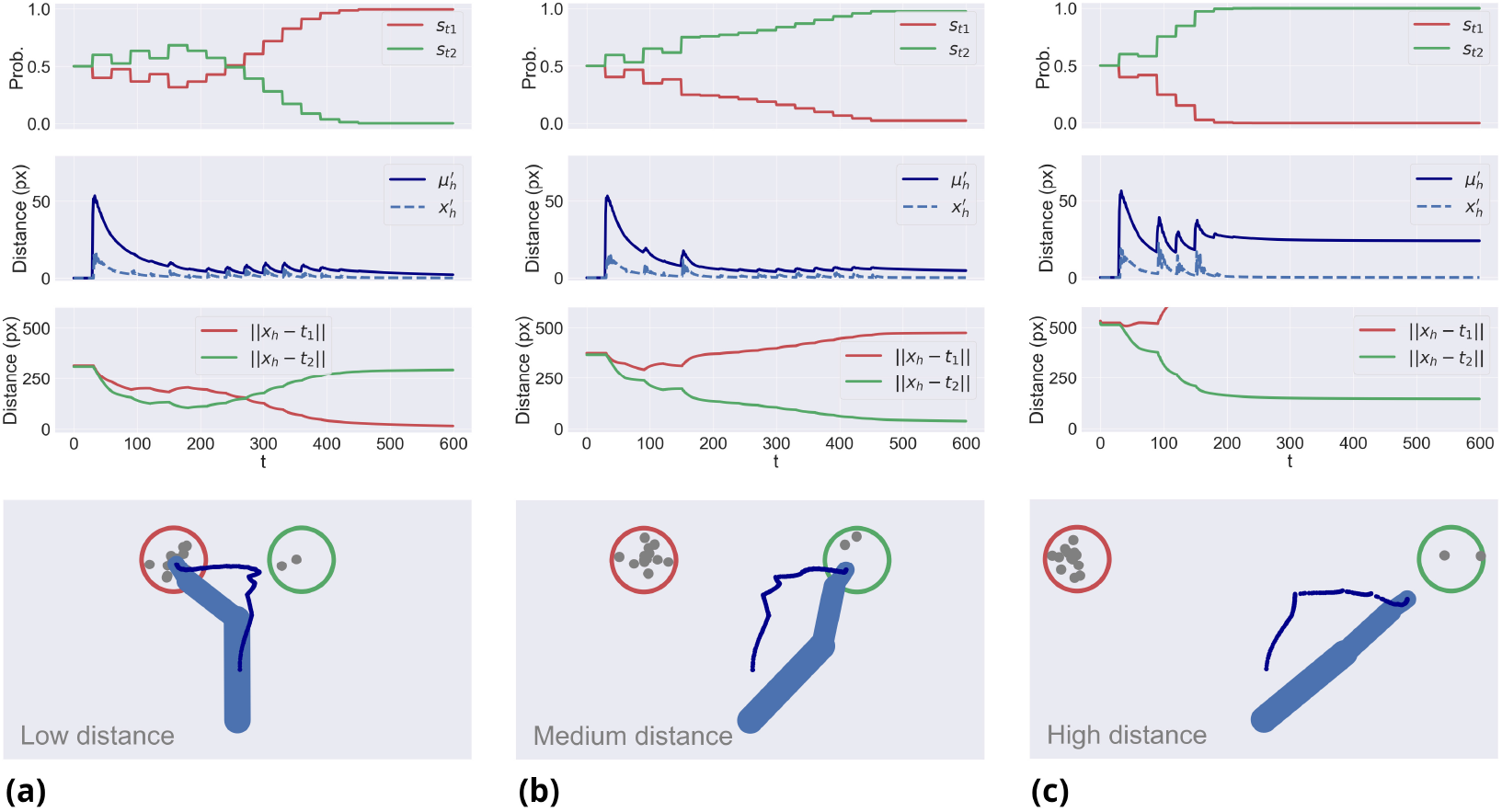
Commitment during an incongruent trial, with (a) low, (b) medium, and (c) high distance between the two targets, with *k*_*h*_ = 0.15 and *α*_*h*_ = 1.0. For each condition, the first plot shows the discrete hidden states ***s***; the second plot shows the L2-norms of the estimated and true hand velocities 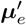 and 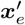; the third plot shows the distances between the hand and the two targets. For each condition, the second row also shows the agent’s final trajectories, in dark blue.

**Fig 6.**
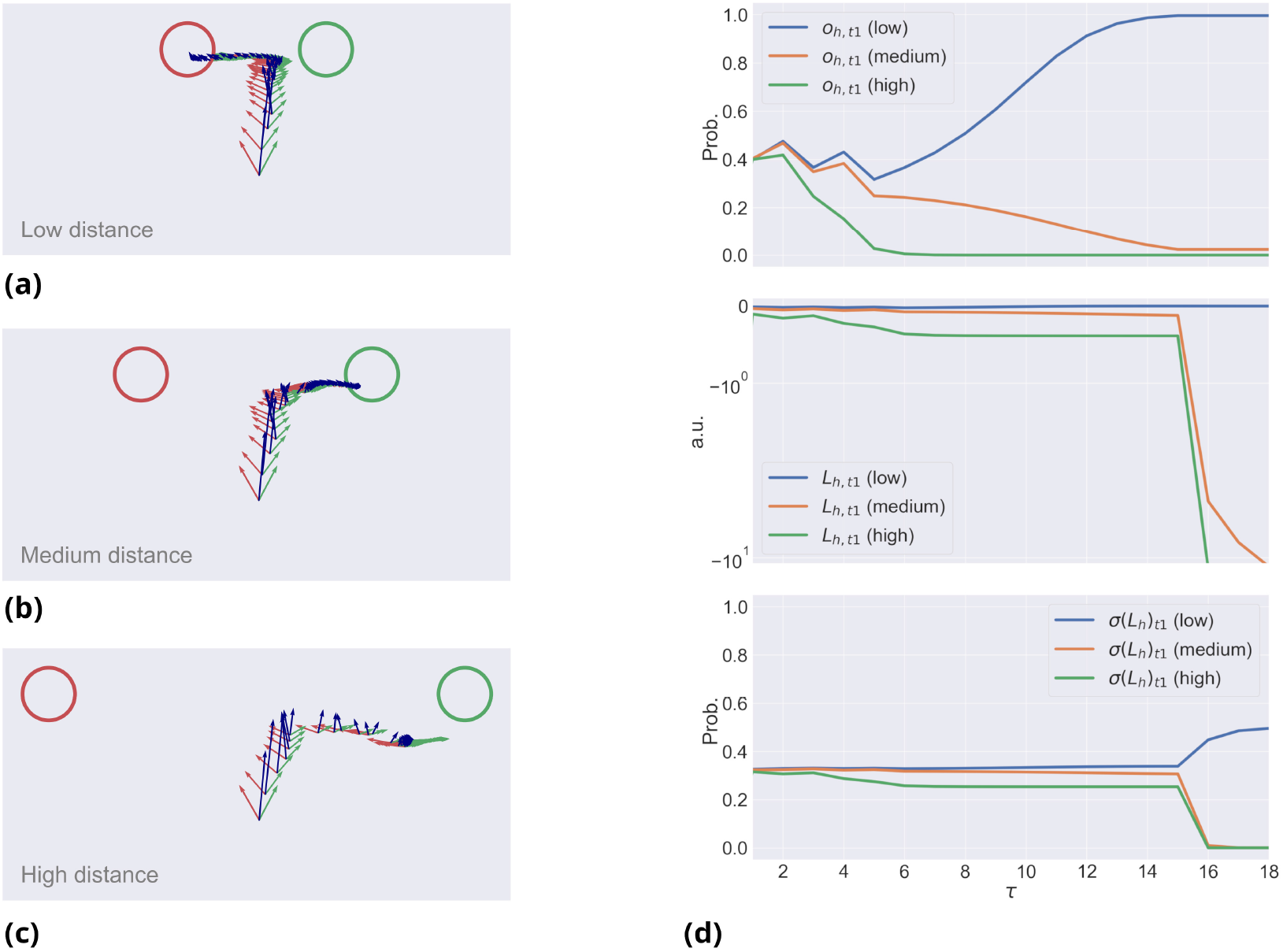
Commitment during an incongruent trial, with (a) low, (b) medium, and (c) high distance between the two targets; continued from Fig 5. (a-c) The panels show the direction and magnitude of the estimated velocity 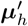 (in dark blue), the potential trajectory needed to reach the first target ***f***_*t*1_ (in red) and the second target ***f***_*t*2_ (in green), for each condition. These potential trajectories are used to estimate which of the two targets is more likely to have generated the current trajectory. (d) The top panel shows the discrete hand dynamics *o*_*h,t*1_ of the first target over discrete time *τ*; the middle panel shows the log evidence ℒ_*h,t*1_ of the hand trajectory associated with the first target, on a logarithmic scale; the bottom panel shows the normalized log evidence of the first target.

**Fig 7.**
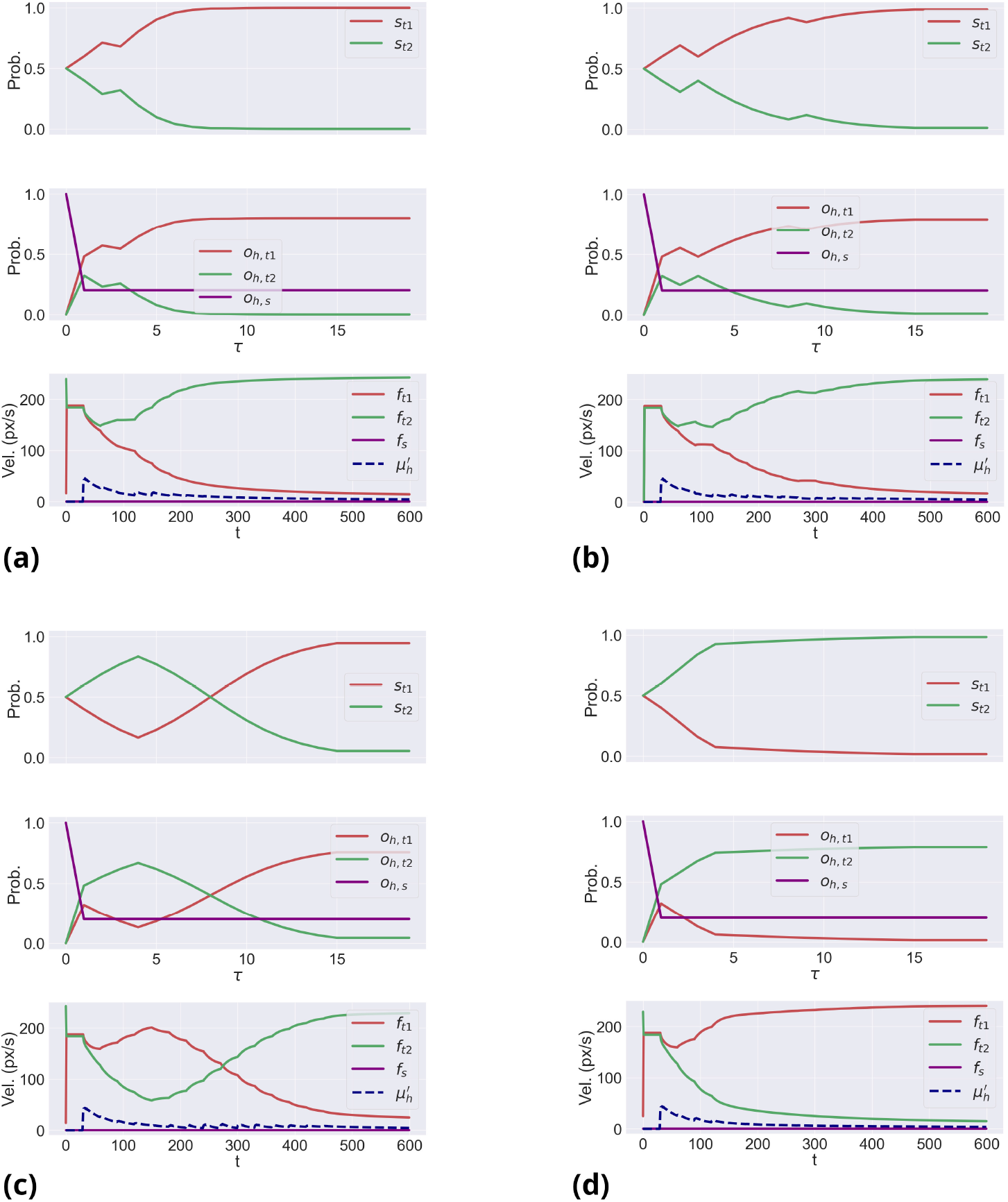
Commitment to an initially selected target resulting from motor inference. Panels a-b compare two agents – without motor inference (a, *k*_*h*_ = 0) and with motor inference (b, *k*_*h*_ = 0.2) – during a *congruent* trial. Panels c-d compare the same two agents – without motor inference (c, *k*_*h*_ = 0) and with motor inference (d, *k*_*h*_ = 0.2) – during an *incongruent* trial. In all conditions, *α*_*h*_ = 0.8. For each panel, the first plot shows the discrete hidden states ***s***; the second plot shows the discrete hand dynamics for reaching both targets and staying in position; the third plot shows the L2-norms of the estimated hand velocity 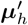, compared with the three potential trajectories ***f***_*t*1_, ***f***_*t*2_, and ***f***_*s*_. Note that although the *stay* dynamics function ***f***_*s*_ is (initially) the closest to the actual trajectory, the related probability *o*_*h,s*_ decreases rapidly, as soon as the predictions ***A***_*h*_***s*** shift toward one of the targets, showing the top-down influence from choice to movement. The fourth (bottom) plot shows the agent’s movement trajectories, in dark blue.

Finally, we considered the tradeoffs between various models: a serial, *decide-only* model that makes decisions between targets by considering the discrete hidden states ***s*** and then moves instantaneously to the selected target; a serial *decide-then-act* model that is identical to the decide-only model, but requires a fixed movement time of 125 time steps (i.e., the average movement time in our setup) to reach the selected target; and two embodied choice models, one with motor inference (*k*_*h*_ = 0.1) and one without it (*k*_*h*_ = 0). For this comparison, we simulated 100 *neutral* trials, by varying the urgency *α*_*h*_ and the drift rate *α*_*c*_ to obtain a wide range of solutions. Fig. 8a shows the results of the comparison. As expected, the speed-accuracy curve of the *decide-only model* appears to be the best (i.e., the leftmost). However, this advantage is misleading, as the model moves instantaneously. It is therefore inappropriate as a model of human decision-making behavior, but it is shown here to illustrate idealized performance. When movement time is accounted for, the *decide-only model* becomes the *decide-then-act model*, which is outperformed by both embodied models – whether without or with motor inference – the latter being the best overall. Furthermore, Fig 8b shows that across various levels of urgency *α*_*h*_, there is a significant correlation between the speed of embodied choice and confidence (i.e., probability of the correct choice) as reported empirically [46]. In sum, embodied choice models show a better speed-accuracy curve than serial *decide-then-act* models. The fact that they afford faster decisions might be particularly advantageous in ecological settings, where slow decisions imply missing valuable alternatives [4, 21]. The relative advantages of including or not including motor inference are likely to be task-dependent. The embodied model using motor inference appears to be advantageous when processing congruent (Fig.7) and neutral trials (Fig.8a), but its commitment to the initially selected target can lead to more incorrect decisions when processing incongruent trials (Fig. 5).

**Fig 8.**
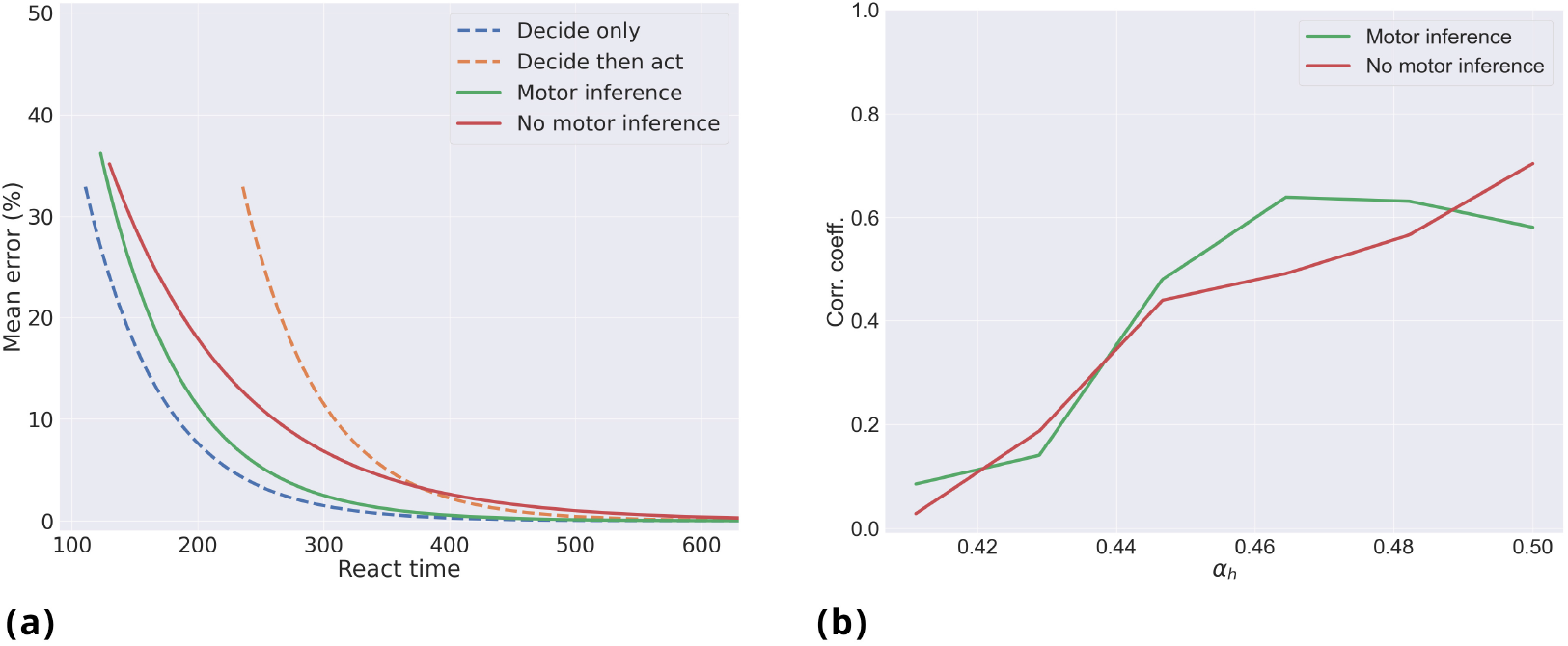
(a) Speed-accuracy curves for four models: two serial models (decide-only, decide-then-act) and two active inference models (with and without motor inference). The former two were sampled by varying the decision threshold in 500 trials, while the latter two were sampled by running 100 trials for different values of urgency (from medium-high urgency, i.e., *α*_*h*_ = 0.7, to low urgency, i.e., *α*_*h*_ = 0.5). The samples were then fitted into a curve. Motor inference was realized with *k*_*h*_ = 0.1. For all conditions, *α*_*c*_ = 0.6. To allow the agents to complete the trials on time with low levels of urgency, we set the maximum trial duration to 630 time steps. (b) Pearson product-moment correlation coefficient between the belief over the hand trajectory 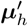 and the probability of the correct choice, computed for two conditions (with and without motor inference), with 500 neutral trials per condition, and with different levels of urgency.

### 2.4 The fourth process: statistical learning and habit formation

During various cognitive tasks such as the Flanker [47] and the Posner task [48], it is possible to learn statistical regularities, such as the probability of the correct response or the validity of cues across trials. In these tasks, trial sequence effects are often reported, indicating that participants form expectations across trials that influence their subsequent responses and movements [49]. The fourth process of our model implements this kind of statistical learning, which simply amounts to keeping count of the Dirichlet priors over the discrete hidden states ***s*** across trials. After every trial, these counts are updated according to:

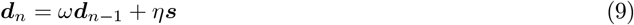

where *n* is the trial number, *ω* is a forgetting factor of older trials, and *η* is the learning rate of new trials (usually initialized with a reasonably high value, reflecting a high confidence over the prior belief, ***d***_0_). Then, the counts are normalized to compute the priors of the correct response for the next trial.

Fig 9 shows the effects of learning the prior over the correct response, during 50 incongruent trials. During the first 10 trials, the correct (left) response remains stable and then it is reversed. During the early trials of the first (learning) phase (dark blue in Fig 9a), the agent moves toward the wrong direction and then changes mind. However, in later trials (dark red in Fig 9a) it gradually begins to move early toward the correct target, anticipating the transition in the accumulation of wrong cues. In parallel, movement onset decreases (Fig 9b). These results show that a strong prior can overcome conflicting evidence. After the reversal at trial 10, the discrete prior for the first target slowly decreases, as the Dirichlet counts for the second target begin to increase (Fig 9b). In early trials, movement curvature increases and movement onset is slower, as the agent is uncertain about the correct distribution underneath the cue sampling. In late trials, movement curvature decreases and movement onset fastens, as the agent learns the novel contingencies.

**Fig 9.**
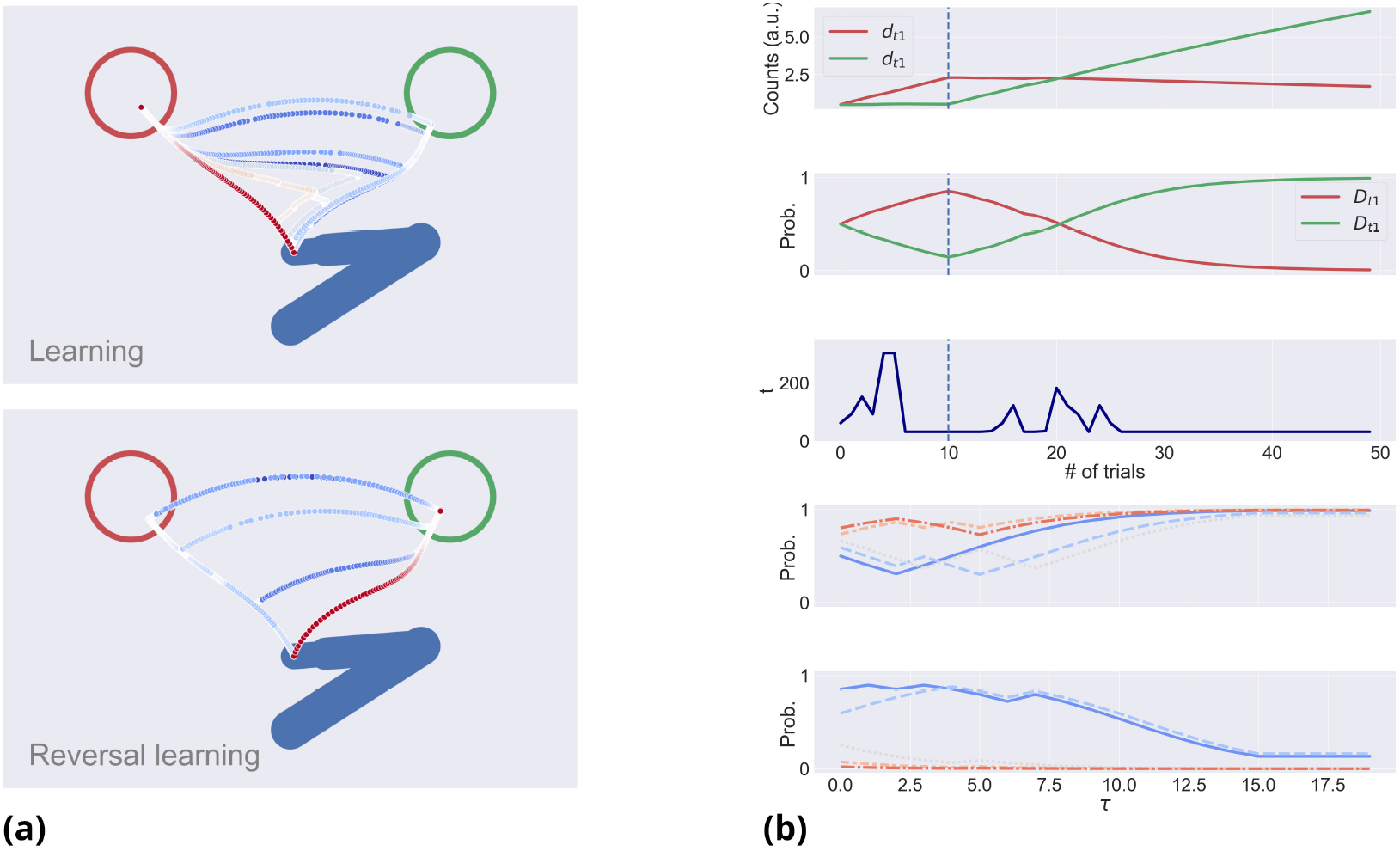
Statistical learning of the prior for the correct choice over 50 incongruent trials. The correct choice is fixed in the first 10 trials, but is reversed in the next 40 trials. (a) Hand trajectories in equally spaced trials, during learning (top) and reversal learning (bottom). Dark blue trajectories represent early trials, while dark red trajectories are late trials. Here, *k*_*h*_ = 0, *ω* = 0.99, *η* = 0.2, *α*_*c*_ = 0.6, and *α*_*h*_ = 0.6. (b) The five panels show Dirichlet counts ***d***; discrete priors ***D***; time step of movement onset across trials; discrete hidden state *s*_1_ in 5 equally spaced trials for learning; and for reversal learning over discrete time *τ*. The vertical dashed lines indicate the time step when reversal occurs.

These results show that our model can incorporate sequence effects that emerge during cognitive tasks [49]. Note that while we focus on learning the prior probability of the correct choice, other model parameters, such as the uncertainties of the likelihood matrices ***A***_*c*_ and ***A***_*h*_, could be updated using the same approach – see also [50].

## 3 Discussion

For many years, the dominant view regarding human and animal behaviors has been that of a serial, *decide-then-act* strategy. However, various studies show that during embodied decisions that require simultaneously specifying and selecting between alternative action plans, the serial view is insufficient. These studies report early movement onset, changes of mind, and the influence of motor costs over decisions, suggesting that decision and action processes unfold in parallel and reciprocally influence each other [10, 12, 13, 21, 27]. Here, we show that these signatures of embodied decisions emerge naturally in active inference: a framework that jointly optimizes decisions and actions under a free energy minimization imperative [29, 31]. Our simulations highlight that four model processes – evidence accumulation, motor planning, motor inference, and statistical learning – form a closed loop, allowing decision and action processes to influence each other reciprocally. The resulting embodied models attain a better speed-accuracy tradeoff, compared to serial models (Fig 8a) – suggesting that they could confer ecological advantages [4, 21].

An innovative aspect of our model is the reciprocal interaction between motor planning and motor inference. During motor planning, the agent’s inference of the correct choice generates predictions about the next discrete hand dynamics, which are converted into a continuous motor plan to reach the associated target. In turn, during motor inference, the agent uses the action dynamics as evidence for the correct choice. In other words, the agent treats its own behavior as a source of information [51]. This mechanism implies that movement stabilizes decisions and creates commitment. Furthermore, it explains key aspects of embodied decisions, such as the fact that motor costs apparent before a task [22] or changing during it [26] influence decision outcomes.

Note that there are two alternative perspectives on (or interpretations of) motor costs in embodied decisions. According to the value-based perspective [21, 25, 27], during movement the agent continuously estimates the cost of the actions to make the alternative choices, then combines the estimated cost with the correct choice (inferred by sensory evidence) to decide the next move. Instead, the perspective offered here is based on (active) inference. Our model does not explicitly compute motor costs, but rather the probabilities of discrete hand dynamics ***o***_*h*_: in short, the agent just tries to infer the correct target from the information at its disposal (including its own movements), and a high motor cost only means a potential dynamics that poorly explains the present context. The two perspectives, active inference and value-based, are mathematically related given the duality of inferential formulations (that use probabilities) and control formulations (that use costs) [52, 53]. But since in active inference (motor) costs are absorbed into (prior) probabilities, its appeal lies in affording embodied decisions using the same inferential machinery required for action control, without the need to compute additional quantities (such as motor costs) on the fly. The same inferential machinery also permits considering other types of prior preferences (or motor costs). For example, including in the model a prior preference for biomechanically simpler movements would automatically favor targets that are closer or can be reached more easily, as empirically observed [22]. Furthermore, while this study focused on perceptual decision tasks, the same model could also be applied to value-based decisions, where different choice targets are associated with varying values or rewards. This could be achieved by assigning a stronger prior preference to targets with higher economic value. Such an extension would also enable the study of how economic value influences motivation and movement vigor in embodied choice tasks [54]. Another way prior preferences (over policies or outcomes) could be leveraged is to account for biases, such as repetition biases that develop across trials. Studies have demonstrated sequence effects in perceptual [55] and visuomotor tasks [56], showing that recent trial history can bias subsequent decisions. The simulations in Fig. 9 illustrate how certain biases or habits can emerge from the statistical learning of previous responses. However, habits can also arise from other aspects of interaction beyond simple repetition. For instance, a reward could become associated with a target and influence movement even if the target changes position – an effect that cannot be explained solely by repeating the same movement. Accounting for these and other cognitive aspects of biases, beyond mere movement repetition, may require increasing prior preferences for abstract outcomes (e.g., reaching a rewarding target regardless of its spatial position). Investigating this possibility remains an open question for future research.

Another key aspect of our model is the fact that by modulating the urgency to move, it can model a range of strategies – from riskier to more conservative – observed in empirical studies [42]. Future work might explore how the generative model advanced here could be inverted, to identify personalized parameters (e.g., a person’s urgency) from behavioral data. While we covered several important aspects of embodied decisions in active inference, we kept the focus on the relationship between discrete decisions and continuous dynamics. A more realistic model would also consider discrete dynamics, which may be crucial for explaining how humans optimize the number of successes accumulated in a limited period – as analyzed in [34]. Furthermore, while our model explains how movement can stabilize decisions, it does not include other stabilization mechanisms, such as the modulation of sensory precision (i.e., Kalman gain) [57–59], which might be covered in future studies.

An important avenue for future research is the empirical validation of the embodied choice models introduced here. In this study, we provided normative arguments for the benefits of embodied models in terms of better speed-accuracy curves compared to serial strategies. Furthermore, we have shown that the model reproduces qualitative aspects of data, such as the increase in the curvature of trajectories and the frequency of “changes of mind” with choice uncertainty [15–18], as well as the correlation between the speed of embodied choice levels and confidence (i.e., probability of the correct choice) as reported empirically [46]. Future studies could use embodied choice models to fit empirical data, by exploiting the fact that different kinds of behavior can be elicited by changing the precision of various likelihood mappings – as shown in [50]. In principle, it would be possible to optimize the precisions of a generative model of a given experimental paradigm to match the behavior of an experimental subject. This would provide an opportunity for computational phenotyping of subjects (or patients) in terms of the (precision of) key components of their generative models [60]. This kind of computational phenotyping has been used to characterize the prior precisions and preferences of psychiatric subjects using choice behavior and, in principle, could be extended to cover the embodied decision-making considered in this work. This would allow for the systematic study of the sensitive dependence of behavior on the relative precision of various likelihood mappings as a potentially important aspect of embodied decision-making. Notably, while each of the four processes illustrated in this study operates in a relatively straightforward manner, their interactions are more complex. It remains an open question whether certain parameter combinations are particularly effective in specific embodied contexts, such as those characterized by varying levels of urgency or precision. For instance, our simulations suggest that the benefits of motor inference depend on both trial statistics (e.g., the proportion of congruent, neutral, and incongruent trials) and urgency, yet the interplay between these factors remains to be explored. Additionally, it is unclear whether individuals can flexibly select the most appropriate set of parameters for each context or whether they instead reuse suboptimal parameterizations across different contexts.

From a neural perspective, embodied choice models align with sensorimotor theories of decision-making and neural evidence from areas such as the posterior parietal cortex (PPC) and dorsal premotor cortex (PMd), where action plans are dynamically represented and compete for selection [61–63]. Importantly, in the embodied choice model, decision-making is a distributed process that integrates multiple (sub)processes or pathways, consistent with the concept of decision-making as a “distributed consensus” rather than a centralized process [28]. The hybrid architecture shown in Fig. 1b could correspond to a (hierarchical) neural architecture involving recurrent dynamics and extensive cortico-basal ganglia-thalamo-cortical loops, which support the continuous interaction between perception, action, and decision processes [64]. For a more detailed discussion of the biological underpinnings of movement control in active inference, see also [65]. Furthermore, the model’s precision-weighting mechanism, which determines how likelihood mappings influence both decisions and movements, could correspond to neuromodulatory processes regulating synaptic efficacy, such as those mediated by dopamine and noradrenaline [66]. Future studies using neuroimaging and electrophysiology could further investigate the neural implementation of our framework and how the variables in Fig. 1b correspond to neural activity during embodied decision-making tasks.

## 4 Methods

### 4.1 Dynamic reduced models in active inference

Active inference is a computational theory that proposes a unifying paradigm to understand cognitive processing and behavior in living organisms. It is based upon the *free energy principle* which states that, in order to survive, every creature must actively minimize surprise [29, 31, 67].

Active inference models have been formulated both in discrete time and in continuous time. However, it is possible to argue that a comprehensive account of living organisms might require *hybrid* models, which combine both discrete and continuous time formulations [68, 69]. For example, in the human nervous system, the cerebral cortex might operate in a discrete state-space, while the low-level sensorimotor loops might be better understood in terms of continuous representations; the interface between the two is attributed to subcortical structures such as the thalamus or the superior colliculus [70]. Hybrid models have been used to simulate many scenarios such as pictographic reading [68], movements under neurological disorders [**?**], or interoceptive control [71]. Here, we briefly present a particular instance of such models useful to infer and act upon dynamic trajectories [36, 37, 45]. See [**?**] for more details about active inference and the free energy principle, and [72] about Bayesian model selection.

Discrete models share resemblances with Hidden Markov Models (HMMs) and are defined as partially observable Markov decision processes (POMDPs) [73, 74]. In particular, they are related to a subfield of machine learning known as *planning as inference* [75, 76]. *Discrete models assume that organisms perceive the environment by optimizing an internal generative model, inferring how external causes lead to sensory signals within the environment (called generative process). Denoting by* ***s*** the discrete hidden states, by ***o*** the discrete outcomes, by ***π*** the policies (which in active inference are sequences of actions), the agent’s generative model after a discrete period *T* is factorized as:

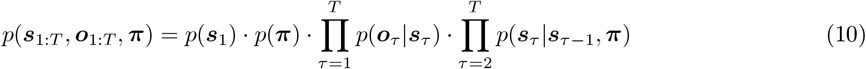

where every distribution is assumed to be categorical:

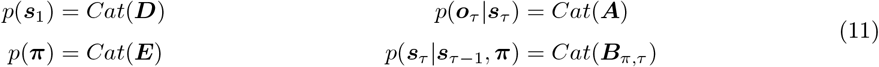

Here, ***D*** is the prior over the initial state, ***E*** is the prior over policies, ***A*** is the likelihood (or observation) matrix, and ***B***_*π,τ*_ is the transition matrix. In order to find the causes of their perceptions, organisms try to infer the posterior distribution:

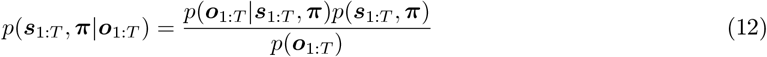

However, this requires computing the intractable model evidence *p*(***o***_1:*T*_). Hence, organisms are supposed to implement some sort of approximate Bayesian inference by relying on an approximate posterior distribution *q*(***s***_1:*T*_, ***π***), and then minimizing the Kullback-Leibler (KL) divergence between the approximate and real posteriors:

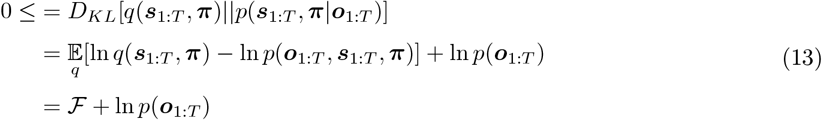

The KL divergence still requires the computation of the log evidence *p*(***o***_1:*T*_); however, we can exploit its non-negativity to instead minimize the first RHS term of the second line of Eq 13 – called variational free energy (VFE) and known in machine learning as the negative *ELBO* – which ensures that surprise − ln *p*(***o***_1:*T*_) is minimized. Then, expressing the approximate posterior by its sufficient statistics ***s***_*π,τ*_ and conditioning upon a specific policy:

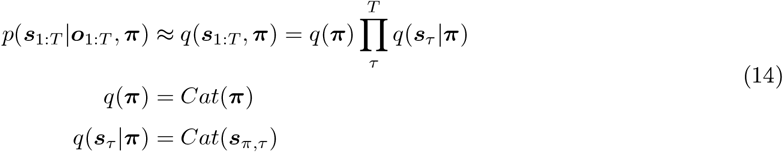

We can infer the most likely discrete hidden state at time *τ* and under policy *π* by computing the gradient of the VFE ℱ_*π*_ of that policy and applying a softmax function to ensure that it is a proper probability distribution:

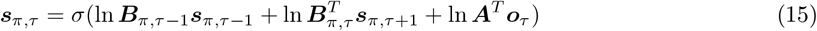

Via VFE minimization, organisms are able to capture the optimal representation of the environment; however, they are unable to perform any sort of future planning. To do this, they additionally consider unobserved outcomes as random variables, and they infer the most likely policy or sequence of actions that will lead to their preferred outcomes. More formally, if we denote by *p*(***o***_*τ*_ |***C***) the probability distribution encoding the agent’s preferred outcomes at some future point *τ*, the optimal policy is found by minimizing the free energy that the agent expects to perceive in the future – called *expected free energy* (EFE) and denoted by 𝒢:

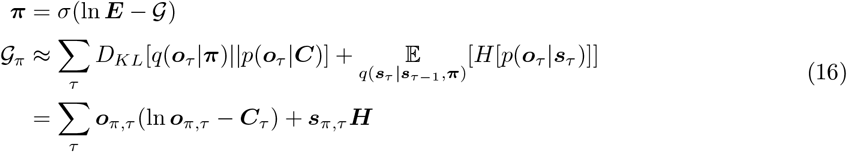

where:

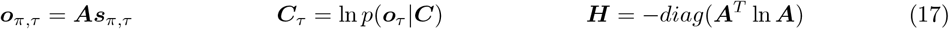

Eq 16 underlies a stark difference with theories of optimal control and reinforcement learning. These theories assume that hidden states have intrinsic values and that agents infer the optimal policy that maximizes the accumulation of future rewards obtained from the environment. Rather, active inference only assumes that each organism believes that the environment will evolve in a specific way, determined by its phenotype. Under this view, action is just another way, complementary to perception, to minimize free energy (hence, surprise), i.e., to reduce the difference between the agent’s prior beliefs and the actual generative process. In other words, by acting, they make future observations coherent with their internal model – a process known as *self-evidencing*. Furthermore, the two components of the second and third lines of Eq 16 entail the well-known tradeoff between exploitation and exploration, with the addition that the latter, also called *ambiguity*, is involved in itinerant and novelty-seeking behavior.

To get a discrete model working with the richness of the continuous input, it is linked to an active inference continuous model [68, 69]. The latter is highly similar to the discrete model described above, with the difference that now only the VFE is minimized (for this reason, a continuous model alone is not capable of advanced decision-making). The standard way to link the two models is by letting a discrete outcome generate a *causal variable* (e.g., the position of a target to reach) that in turn produces a continuous trajectory; in this way, inverting the model means to infer the target based on the continuous trajectory, and then finding the discrete outcome that best explains the inferred target by comparing it with some fixed positions that the agent knows a-priori. In order to render the agent’s representation more flexible, we can instead generate a continuous trajectory directly by a discrete outcome. Here, we briefly describe this alternative approach.

First, we model the continuous environment with the following non-linear stochastic equations:

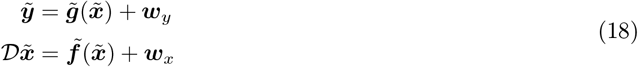

where 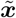 are continuous hidden states, 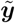 are continuous observations, 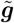 is a likelihood function defining how hidden states cause observations, 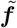 is a dynamics function expressing how the hidden states evolve, and the letter *w* indicates Gaussian noise terms. Note that the symbol ~ denotes generalized coordinates of motion encoding instantaneous trajectories (e.g., position, velocity, acceleration, and so on), which in a continuous formulation replace the future states. Also, D is a differential operator that shifts every coordinate of motion by one. The joint distribution of this *hybrid* generative model is factorized into the following:

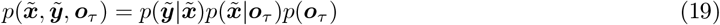

where the first two distributions are assumed to be Gaussian, while the last one is the categorical (likelihood) distribution defined before:

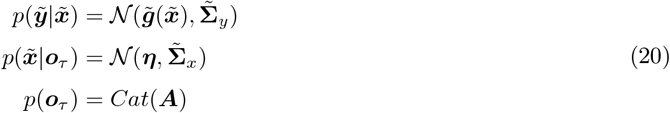

Here, ***η*** is the mean (also called *full prior*) of a complex model that defines the actual evolution of the hidden states. We suppose that the agent maintains *M* probability distributions which are reduced versions of this full model. Each reduced distribution corresponds to a particular *discrete dynamics o*_*τ,m*_ and encodes a specific way that the agent thinks the environment may evolve [36, 37]:

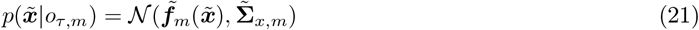

where 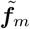 is the dynamics function of the *m*th reduced model, and 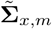 the related precision. These alternative hypotheses act as empirical priors at the lower continuous level. Following the theory of Bayesian model reduction [72, 77], in order to infer the posterior of the full model we first have to compute the posterior probability of each reduced model. In fact, reduced means that the likelihood of some data is equal to that of the full model and the only difference rests upon the specification of the priors – hence, the posterior of a reduced model can be expressed in terms of the posterior of the full model:

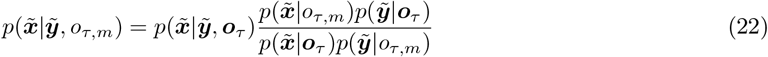

As in the previous discrete formulation, we introduce a full and M approximate posteriors, here assumed to be Gaussian:

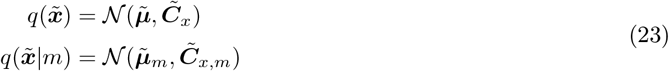

where 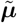 and 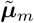 are the full and reduced posterior means (also called *beliefs*), while 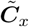 and 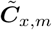 are the full and reduced posterior precisions. Replacing the real posteriors with these approximate posteriors, we can write the free energy of each reduced model in terms of the full model. As before, maximizing each reduced free energy makes it approximate the log evidence:

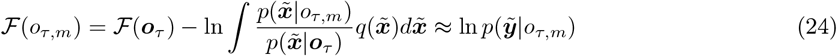

This relation implies that the free energy related to each discrete dynamics *o*_*τ,m*_ can be found from the approximate posterior 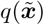 of the full model, without directly computing the reduced posteriors. With the Gaussian approximation of Eq 23, the *m*th reduced free energy breaks down to a simple formula:

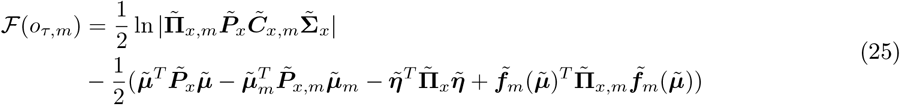

where the mean and precision of *m*th reduced model are:

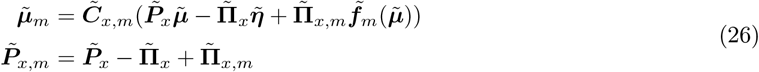

and we wrote the covariances 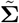 and 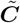 in terms of precisions 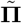 and 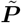. In short, the free energy assigns a score to each reduced model based on its fit with the full posterior, and two models *i* and *j* can be compared through a log-Bayes factor ℱ(*o*_*τ,i*_) − ℱ(*o*_*τ,j*_). Crucially, this posterior is a continuous trajectory and the agent’s hypotheses are constantly updated via sensory observations. If a discrete step *τ* corresponds to a continuous period *T*, we integrate the reduced free energies over this period and compare them with the prior surprise − ln ***o***_*τ*_. This results in an ascending message that infers the most likely discrete dynamics ***o***_*τ*_ that may have generated the current continuous observations:

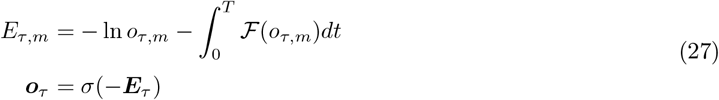

Instead, the descending message is computed as a simple Bayesian model average of the agent’s hypotheses, e.g., by weighting the discrete dynamics *o*_*τ,m*_ of a reduced model with the related dynamics function 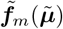:

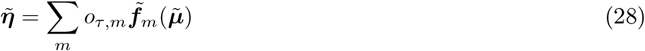

Then, 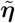 embeds a prior over a continuous trajectory, which steers the inference of the continuous hidden states toward preferred outcomes. The posterior over the hidden states is finally found by computing the gradient of the free energy of the full model with respect to the mean 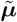 of the full approximate posterior, and updating the latter via gradient descent:

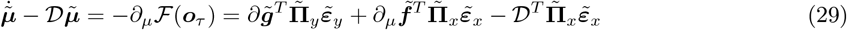

This update is expressed in terms of message passing of the following prediction errors:

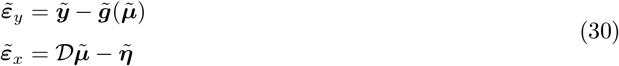

and has a specular form of the discrete update of Eq 15, except that now the inference is done over a continuous path – hence the additional term 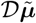. This minimization entails the process of perception, i.e., conforms the agent’s beliefs to the perceived sensations. In order to conform the environment to the agent’s beliefs (in short, to act), the free energy can be minimized with respect to the motor commands ***a***. This additional mechanism reduces to a minimization of sensory prediction errors:

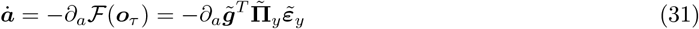

where 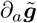 is an inverse mapping from sensations to actions. This mapping implements a dynamics inversion and is thought to be realized by classical reflex arcs in the spinal cord [78, 79].

### 4.2 Inverse kinematics in active inference

Several methods have been proposed on how to realize inverse kinematics in active inference. The most common approach for simulating a reaching task is to encode a target location in extrinsic coordinates as a causal variable of a continuous model. This variable generates a prediction for the 1st-order (e.g., velocity) of the hidden states – encoding the agent’s joint angles in intrinsic coordinates – which therefore act as a dynamic attractor toward the desired location. In this simple representation, the link between the causal variable and the hidden states performs an inverse kinematics of the target location, whose product (i.e., a possible agent’s configuration with the hand at the target) is compared with the extrinsic position of the hand computed via forward kinematics. In this simple representation, both (intrinsic and extrinsic) reference frames are used in a single active inference level.

An alternative and more powerful approach exploits an aspect of the theory inherited from predictive coding, i.e., that the nervous system manages to approximate the real posterior by building a hierarchical architecture wherein a particular level acts as an observation for the level above and as a prior for the level below [43, 44]. In this way, higher levels can construct increasingly richer and more invariant representations of the environment, similar to deep generative models of neural networks. Since a level communicates only with the levels immediately above and below, the overall generative model can be factorized into independent distributions, and every level can be analyzed as in the general formulations of discrete and continuous active inference. Specifically, the hierarchical approach to inverse kinematics is to design a structure of two levels, wherein an *intrinsic unit* encoding the agent’s joint angles has a causal influence on an *extrinsic unit* encoding the agent’s hand. This approach follows the natural flow of the generative process, with an efficient decomposition between intrinsic and extrinsic dynamics. Here, we briefly describe this second approach.

The higher-level intrinsic unit 𝒰_*i*_ is governed by the following equations:

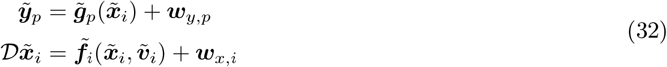

where 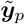 are generalized (e.g., position, velocity, and so on) proprioceptive observations, 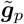 are the generalized proprioceptive likelihoods, 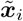 are intrinsic hidden states encoding the instantaneous trajectory of the agent (e.g., every joint angle of the arm), 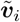 are intrinsic causal variables encoding the trajectory priors, and 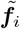 are the related generalized dynamics. Also, the letter *w* indicates Gaussian noises.

Instead, the lower-level extrinsic unit U_*e*_ follows the system:

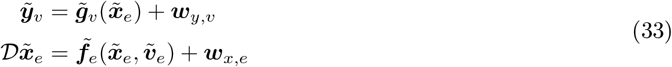

where the letter *v* indicates the visual (or, more in general, exteroceptive) domain. The two units are related via connection between the 0th (i.e., position) temporal orders:

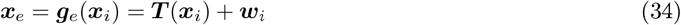

where ***T*** performs a forward kinematics between the agent’s joint angles and its hand. As a consequence, the backward message from 𝒰_*e*_ to 𝒰_*i*_ realizes an inverse kinematics of an *extrinsic prediction error* conveying the difference between the prediction of 𝒰_*i*_ and the actual extrinsic belief of 𝒰_*e*_:

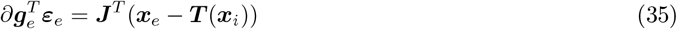

In this way, extrinsic trajectories (e.g., linear or circular motions) are easily realized by defining appropriate dynamics in 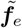 which then travels back to infer the most likely agent’s configuration corresponding to that trajectory. The updates of the belief over intrinsic and extrinsic hidden states are the following:

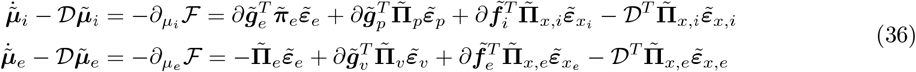

For both units, we note an extrinsic prediction error acting either as a prior or as an observation, a sensory-level observation (either proprioceptive or exteroceptive), and a dynamics prediction error from previous or successive temporal orders. This two-level architecture is highly effective in tasks that require dynamic constraints in both intrinsic and extrinsic domains (e.g., when moving the arm while keeping the hand’s palm up) but fails when simultaneous coordination of multiple limbs is needed. In this case, we can extend the model by designing an *intrinsic-extrinsic module* for every degree of freedom of the agent’s body. Then, a dynamic attractor at the last (e.g., hand) level results in a prediction error that is backpropagated throughout the whole hierarchy, eventually inferring an appropriate hierarchical configuration of the body.

## Code and data

Code and data are available at: https://github.com/priorelli/embodied-decisions

## Acknowledgments

This research received funding from the European Research Council under the Grant Agreement No. 820213 (ThinkAhead) to GP, the Italian National Recovery and Resilience Plan (NRRP), M4C2, funded by the European Union – NextGenerationEU (Project IR0000011, CUP B51E22000150006, “EBRAINS-Italy”; Project PE0000013, “FAIR”; Project PE0000006, “MNESYS”) to GP, the European Union’s Horizon H2020-EIC-FETPROACT-2019 Programme for Research and Innovation under Grant Agreement 951910 to IPS, and the Ministry of University and Research, PRIN PNRR P20224FESY and PRIN 20229Z7M8N. The GEFORCE Quadro RTX6000 and Titan GPU cards used for this research were donated by the NVIDIA Corporation. The funders had no role in study design, data collection and analysis, decision to publish, or preparation of the manuscript. This work has been carried out while MP was enrolled in the Italian National Doctorate on Artificial Intelligence run by Sapienza University of Rome in collaboration with the Institute of Cognitive Sciences and Technologies, National Research Council of Italy. We used a Generative AI model to correct typographical errors and edit language for clarity.

